# Evolving Membrane-Associated Accessory Protein Variants for Improved Adeno-associated Virus Production

**DOI:** 10.1101/2023.07.18.549400

**Authors:** Adam J. Schieferecke, Hyuncheol Lee, Aleysha Chen, Vindhya Kilaru, Justin Krish Williams, David. V. Schaffer

**Author notes:** Denotes co-first author (Adam J. Schieferecke, Hyuncheol Lee). Correspondence should be addressed to D.V.S. and A.J.S.; Corresponding author: David. V. Schaffer. Ph.D., University of California, Berkeley 274 Stanley Hall, Berkeley, CA 94720-3220. All work was performed in Berkeley, California, United States of America.

## Abstract

Manufacturing sufficient Adeno-Associated Virus (AAV) to meet current and projected clinical needs is a significant hurdle to the growing gene therapy industry. The recently discovered membrane-associated accessory protein (MAAP) is encoded by an alternative open reading frame in the AAV *cap* gene that is found in all presently reported natural serotypes. Recent evidence has emerged supporting a functional role of MAAP in AAV egress, though the underlying mechanisms of MAAP function remain unknown. Here, we show that inactivation of MAAP from AAV2 by a single point mutation that is silent in the VP1 ORF (AAV2-ΔMAAP) decreased exosome-associated and secreted vector genome production. We hypothesized that novel MAAP variants could be evolved to increase AAV production and thus subjected a library encoding over 1E6 MAAP protein variants to five rounds of packaging selection into the AAV2-ΔMAAP capsid. Between each successive packaging round, we observed a progressive increase in both overall titer and ratio of secreted vector genomes conferred by the bulk selected MAAP library population. Next-generation sequencing uncovered enriched mutational features, and a resulting selected MAAP variant containing missense mutations and a novel C-terminal domain increased overall GFP transgene packaging in AAV2, AAV6, and AAV9 capsids.

## Introduction

Adeno-associated virus (AAV) is a dependoparvovirus whose natural genome contains ∼4.7 kb of single-stranded DNA that is flanked by inverted terminal repeats (ITRs) and encodes up to ten known proteins in a highly overlapped fashion.^1–3^ The *rep* gene encodes four protein products named according to their molecular weight: Rep78 and Rep68 perform important functions related to cell cycle modulation,^4, 5^ transcriptional activation,^6, 7^ and genomic replication,^8, 9^ whereas Rep52 and Rep40 play essential roles in loading nascent ssDNA genomes into assembled capsids.^10, 11^ Downstream of *rep* lies the *cap* region, which includes ORFs that encode up to six known protein products: VP1, VP2, and VP3 are structural proteins that assemble to form the capsid;^12, 13^ the assembly activating protein (AAP) targets VP proteins to the nucleus and is involved in capsid assembly;^14–16^ the X protein is found in most natural AAV serotypes though its function and underlying mechanisms are yet to be fully elucidated;^17, 18^ and the membrane-associated accessory protein (MAAP) yields a 13 kDa protein product encoded by a +1 open reading frame (ORF) that is found in all presently reported natural serotypes.^3^ In addition, AAV requires the function of helper genes from larger viruses such as adenoviruses or herpesviruses to complete its replication cycle.^1, 19–21^

AAVs offer multiple advantageous properties for use as clinical gene delivery vectors, including the ability to package heterologous genetic payloads enabled by providing natural viral *rep* and *cap* genes *in trans* of an ITR-containing vector genome,^2, 22^ inherent non-pathogenicity and relatively low immunogenicity in humans leading to an extensive clinical safety profile,^23–25^ long-term episomal expression with low risk of genomic integration,^26, 27^ established techniques to produce vectors in cell culture,^28, 29^ physical stability,^30, 31^ efficient transduction of both dividing and non-dividing cells,^32, 33^ and engineerable tropism for specific cell types.^33, 34^ Accordingly, AAVs are the subject of 161 active- or recruiting status clinical trials^25^ and three FDA-approved products as of May, 2023.^35–37^ As the number of clinical-stage AAV products increases, manufacturing the high quantities of good manufacturing practice (GMP)-grade AAV necessary to address current and especially future clinical dosing needs presents a significant challenge.^38, 39^

First reported in 2019, MAAP initiates translation at a CTG codon in the +1 ORF relative to the VP1 ORF,^3, 40^ associates with the plasma membrane,^3, 41^ and increases AAV2 packaging fitness through competitive exclusion as indicated by wild-type AAV2 outcompeting MAAP-null AAV2 in co-infection assays.^3^ Recently, evidence supporting a functional role for MAAP in AAV secretion has emerged, including observations that ablating MAAP reduces AAV8 secretion levels, trans-complementation of AAV8 MAAP can rescue secretion levels across several different AAV serotypes, and AAV8 MAAP interacts with the surface of extracellular vesicles (EVs).^41^ Furthermore, C-terminal MAAP truncations have been shown to confer a modest increase in AAV2 or AAV8 vector genome production.^40–41^ Here, we showed that ablation of MAAP decreased both EV-associated and secreted AAV2 vector genome titer 72 hours post-infection. Furthermore, we postulated that directed evolution, diversification of a target gene followed by selection for improved function, could be utilized to engineer MAAP variants that confer increased recombinant AAV packaging in HEK293 cells. To test this hypothesis, we generated an error-prone PCR library of over 1E6 MAAP variants, which we subjected to five rounds of iterative packaging into an AAV2 capsid using a *cap* gene in which MAAP expression was inactivated (AAV2-ΔMAAP). With each successive packaging round, we observed a progressive increase in both overall titer and ratio of secreted vector genomes conferred by the bulk-selected MAAP library population. Next-generation sequencing revealed mutational features, including novel C-terminal domains. When stably expressed in HEK293 cells, leading selected MAAP variants increased packaging of a GFP transgene into AAV2, AAV6, or AAV9 capsids. Our results contribute new insights into AAV biology, describe a broadly useful directed evolution strategy for engineering non-structural AAV genes, and uncover novel MAAP variants with potential to improve upstream recombinant AAV manufacturing.

## Results

### MAAP is an important factor for AAV2 secretion

It was recently shown that MAAP affects the kinetics and quantity of AAV8 secretion.^41^ To confirm the effect of MAAP on the production of AAV2, we generated an AAV2 genome from which the endogenous expression of MAAP was ablated (AAV2-ΔMAAP) by introducing a point mutation that results in an early stop codon at the 19^th^ leucine of MAAP but is silent in the overlapping VP1 ORF (Fig 1A-1B). Upon packaging into HEK293 cells and sampling at 72 hours post-transfection, we observed a marked decrease in the number of vector genomes of AAV2-ΔMAAP secreted into the supernatant relative to wild-type AAV2 (Figures 1C-1F). Restoration of MAAP expression from a plasmid *in trans* partially rescued the titer of AAV2 secreted in cellular supernatant (Figures 1C-1D). Notably, a statistically significant difference was not observed in the ratio of vector genomes per capsid between wild-type AAV2 vs AAV2-ΔMAAP associated with cells nor supernatant, suggesting that the role MAAP plays in AAV2 production is independent of capsid genome loading (Figures 1E-1G).

**Figure 1.**
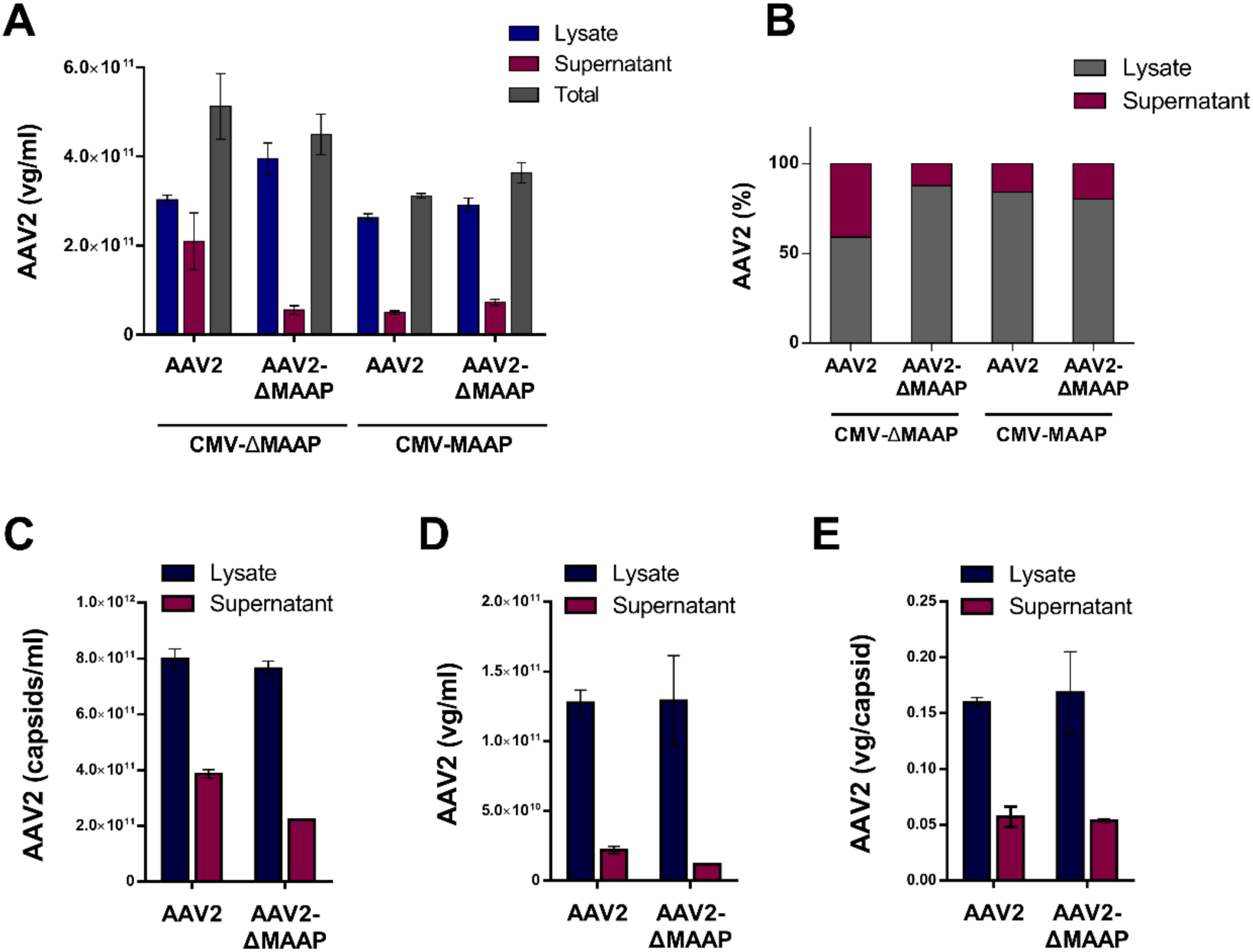
AAV2 MAAP affects viral secretion. A) Knockout of MAAP from AAV2 increases proportion of vector genomes retained in the cell (lysate) 72 hours post-transfection as shown by qPCR. Navy Blue = cell-associated (lysate) AAV2 titer. Maroon = supernatant-associated (secreted) AAV titer. Grey = Total AAV titer. The inactivation of MAAP results in decreased secretion (more AAV retained within the cell). B) Same data as in A showing the ratio of secreted AAV2 (maroon) vs intracellular AAV (grey). Inactivation of MAAP2 results in a decrease in the secretion ratio.Reconstituting MAAP in trans partially restores the secretion of AAV2. C) ELISA of AAV2 capsids. Knockout of MAAP results in a decrease in secretion of AAV2 capsids, corroborating trends observed in vector genome levels. D) Vector genomes are secreted less in the absence of MAAP. E) Vector genome / capsid ratios are unaffected by knockout of MAAP from AAV2. All data are representative of at least two independent experiments. Error bars, mean ± SD.

### Ablation of MAAP decreases AAV2 association with extracellular vesicles (EVs)

To directly analyze the effects of MAAP inactivation on the secreted AAV population, EVs from HEK293 cells transfected with pHelper plus either AAV2 or AAV2-ΔMAAP were isolated into individual fractions, which underwent quantification and molecular profiling via Western Blot staining for EV-associated viral and cellular proteins (Figure 2A-2B). Specifically, following a series of collections and centrifugations of supernatant samples from AAV transfection cultures, the aggregate crude pellet was suspended and floated to the interface between 40% and on a 55% sucrose in a step gradient and then fractionated, wherein the first 5 fractions were stained for protein markers and analyzed for counts of EV particles and EV-associated AAV2 titers (Figure 2A). Notably, lower levels of EVs were secreted from AAV2-ΔMAAP-transfected cell lines compared to cells not producing AAV (Figure 2B-2C). Although cells producing AAV2Δ MAAP showed no significant difference in the levels of EVs produced relative to cells producing AAV2 for which MAAP was intact (Figure 2D), the AAV vector genomes and capsid proteins associated with isolated EV fractions were lower for AAVΔ MAAP vs. wild-type AAV2 samples. These results indicate that MAAP promotes AAV secretion, at least in part, through the association of AAV with EVs (Figures 2E-2F).

**Figure 2.**
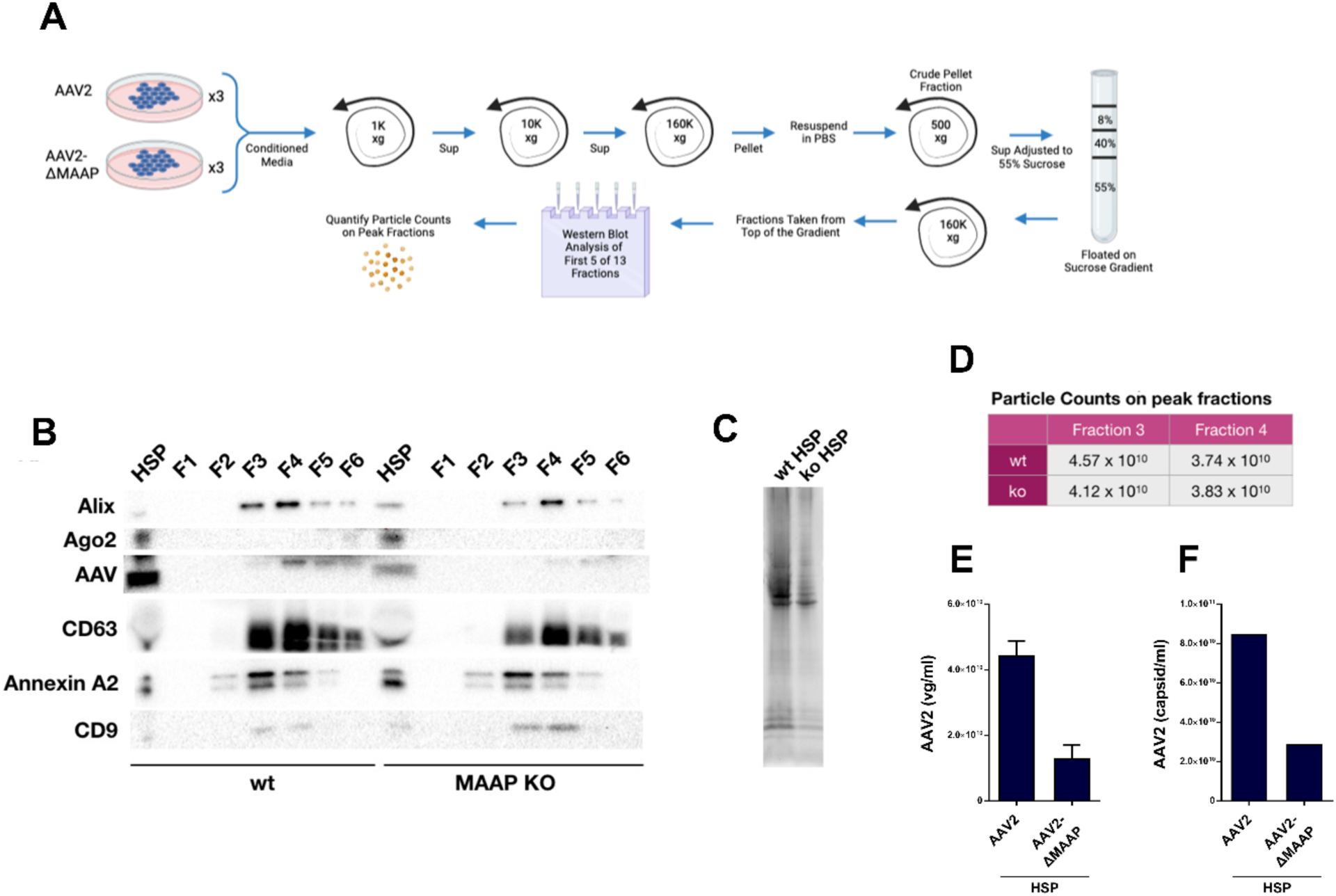
MAAP is required for optimal AAV2 association with exosomes. A-B) AAV-GFP was packaged with AAV2-WT or AAV2-MAAP-null packaging plasmids in HEK293 cells. Cell culture media was changed with exosome-free DMEM at 1 day post transfection and supernatant was harvested at 3 day post transfection for exosome analysis. -Schematic showing the process for isolating and analyzing exosomes. *Created on BioRender.com* (A) and immunoblot of exosome markers and AAV capsid protein in each fraction (B). D) EV particle counts were unaffected by the presence or absence of MAAP. E-F) qPCR-based vector genome titer (E) and capsid ELISA (F) to quantify EV-associated AAV2. Data are representative of two independent experiments (E and F). Error bars, mean ± SD.

### Directed evolution of MAAP results in increased AAV2 production and strong enrichment of unique variants

Directed evolution is an especially powerful approach for engineering proteins for which underlying mechanistic knowledge is insufficient to enable rational design, such as for the recently discovered MAAP. Based on recent evidence reported by other groups^3, 40, 41^ and our own data (Figures 1-2), we hypothesized that directed evolution could yield novel MAAP variants that improve AAV production in industrially relevant cell lines. To test this idea, we generated an error-prone PCR library encoding over 1E6 MAAP protein variants (Figure 4A), which we incorporated downstream of a CMV promoter and flanked by AAV2 ITRs in a plasmid called pMAAP-Library (Figures 3A). The resulting MAAP library was packaged into AAV2 capsids in HEK293 cells by co-transfection with pHelper and a plasmid containing the *rep* and *cap* genes in which endogenous MAAP was inactivated (pAAV2-Δ MAAP). The AAV2-packaged MAAP library was subsequently used for iterative rounds of supernatant-based selection as described in Figure 3A and the methods. Briefly, pMAAP-Library, pHelper, and pAAV2-Δ MAAP were triple transfected into HEK293 cells and incubated for 96 hours; at which point secreted AAV2-ΔMAAP was sampled by removing the cells from the supernatant, titered by qPCR, and infected on to fresh HEK293 cells at MOI = 100 for the subsequent round of packaging with the assistance of co-transfected pHelper and pAAV-ΔMAAP, a process that was iterated four times. The bulk-selected MAAP library population was titered for each successive selection round and showed a progressive and significant increase in both overall titer and the ratio of secreted vector genomes present in the supernatant (from 15.8% to 60.0%) with each round. (Figure 3B-3C).

**Figure 3.**
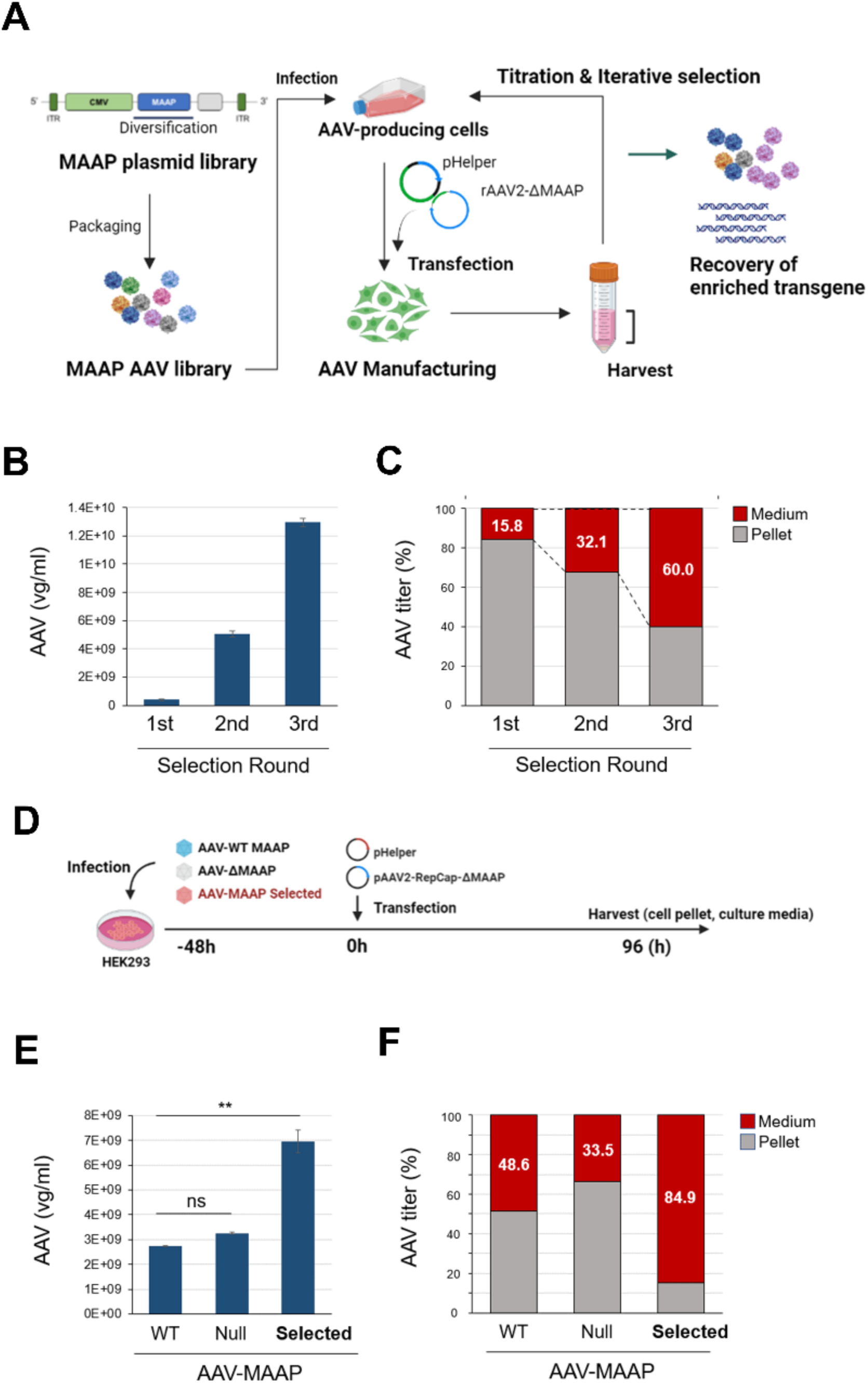
Directed evolution approach for MAAP. A) Schematic of directed evolution approach to generate MAAP variants that confer improved AAV2 production in HEK293 cells. *Created on BioRender.com* **B)** Head-to-head comparison of total vector genome titer (obtained by qPCR) of the MAAP library bulk population after one, two, or three rounds of AAV2-MAAP-null packaging selection relative to the titer after the first round. **C)** Ratio of secreted vector genomes for samples described in B. **D)** Schematic for head-to-head comparison of individual MAAP variants. *Created on BioRender.com.* **E)** Head-to-head comparison of relative total vector genome titer (obtained by qPCR) when packaged into AAV2-MAAP-null containing wild type (WT) MAAP2, no MAAP, or the selected MAAP library bulk population. **F)** Ratio of secreted AAV2 for samples in E. Error bars, mean ± SD.

**Figure 4.**
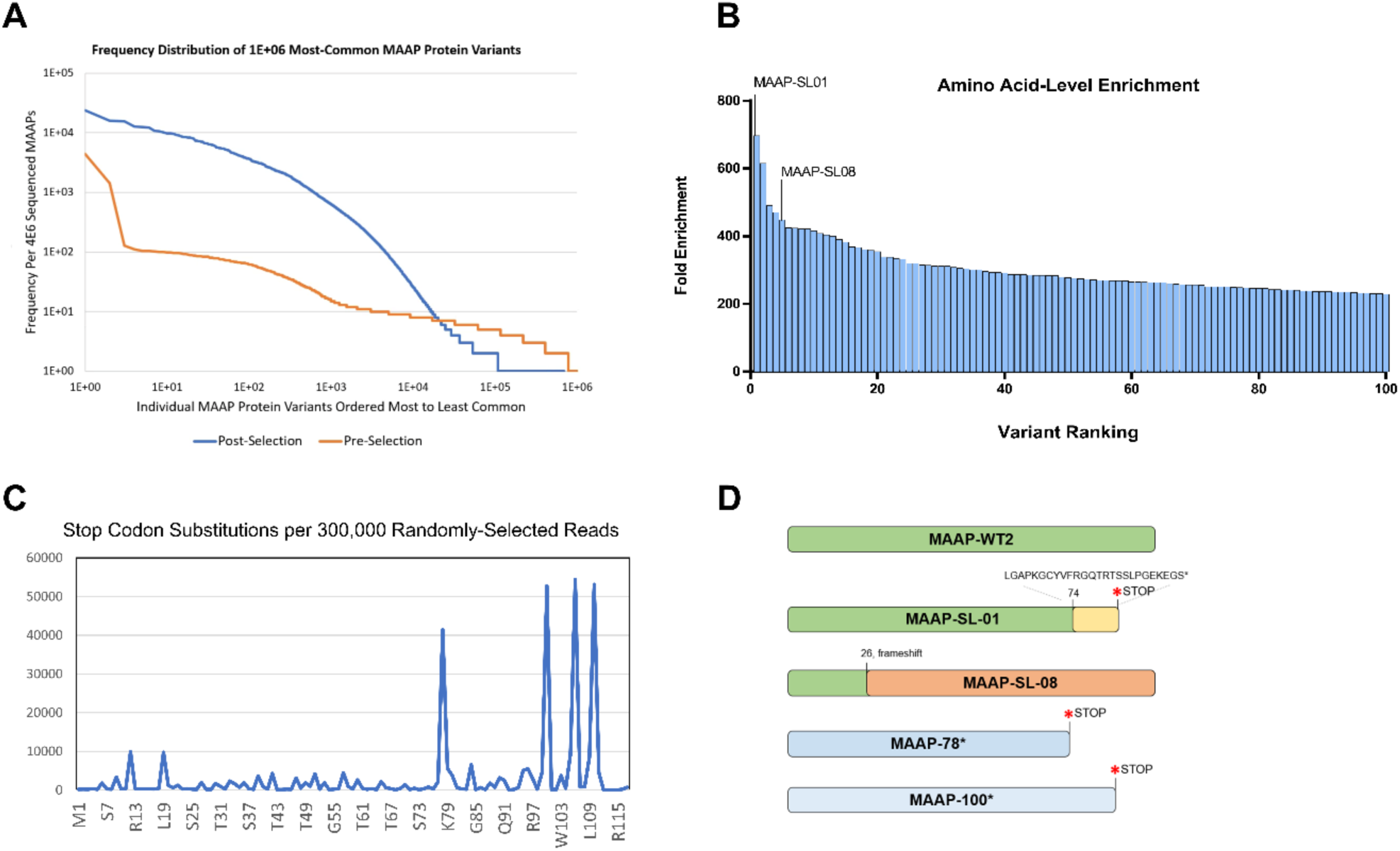
Next-generation sequencing of selected MAAP library uncovers strong enrichment of common features. A) MAAP variants present in the pre-packaging (orange) or post-selection (blue) error-prone MAAP library were aligned, trimmed, translated into amino acid sequence, and ordered based on their frequency. The number of overall MAAP variant sequences with >1 read decreased by approximately one order of magnitude and leading MAAP variants increased in prevalence in the post-selected MAAP library population relative to the pre-packaging MAAP library population. B) Quantitation of fold enrichment of the top 100 unique amino acid sequences present in the post-selection library population relative to the pre-packaging library population. Selected MAAP variants were named based on the order of their enrichment. C) Early stop codons resulting in C-terminal truncations occurred frequently in the post-selected MAAP library population. D) Schematic of MAAP-WT2, MAAP-WT2 truncated MAAP sequences (MAAP-L78* and MAAP-L100*), and two selected leading variants from our directed evolution process (MAAP-SL01 and MAAP-SL08).

Following four selection rounds, we performed a head-to-head comparison of AAV2 production levels conferred by wild-type AAV2 MAAP (MAAP-WT2), empty vector (no MAAP), or the bulk selected MAAP library population, each packaged into AAV2 capsids produced from pAAV2ΔMAAP (Figure 3D). Encouragingly, the bulk selected MAAP library population exhibited over a two-fold increase in overall titer (Figure 3E) and an increase in the relative level of AAV associated with cells vs. supernatant (84.9% AAV found in culture medium, Figure 3F). Next-generation sequencing of the pre- vs post-selected populations showed both a decrease in the overall number of unique MAAP variants and an increase in the frequency of specific variants (Figure 4A). Due to MAAP’s small coding sequence size of 360 nucleic acids, an individual 350 paired-end Illumina read can encompass a full MAAP coding sequence, and the resulting full-length MAAP sequences were translated into amino acid sequence. Individual amino acid sequences were enriched up to 697-fold relative to their prevalence in the pre- selected library population (Figure 4C). Interestingly, we observed strong enrichment of early stop codons in the post-selected MAAP dataset (Figure 4B), a result that corroborates recent findings that truncated MAAP sequences increase secreted AAV8 or AAV2 levels.^40, 41^ However, additional common mutational features, importantly including missense mutations as well as frameshifts that resulted in novel C-terminal domains (Figure 4D, Table S1), were also enriched on the amino acid level (Figure 4C-4D).

### Isolated MAAP variants confer increased recombinant AAV2 packaging

We next determined whether individual MAAP variants enriched in the post-selected population were responsible for the observed increase in AAV2 packaging and secretion (Figure 3). Five MAAP clones encompassing some of the most enriched mutational features observed in the NGS dataset and chosen based on their order of enrichment were selected for head-to-head comparisons of their effect on AAV2 packaging (Figures S3A-S3B). For example, MAAP-SL08 was a fundamentally novel variant that contained the first 26 amino acids of MAAP-WT2 (normally 119 amino acids in length) followed by a frameshift into the VP1 amino acid positions 54-196 with nine missense mutations scattered three to 20 amino acids apart (Figure 4C). By comparison, the most enriched variant in the dataset, SL01, contained the first 73 amino acids of MAAP-WT2 with the exceptions of 13 missense mutations followed by a frameshift mutation that resulted in a novel 26 amino acid-long C-terminal domain (Figure 4D and Table S1). It was recently reported that truncated MAAP sequences containing early stop codons at the 78^th^ or 100^th^ amino acid positions but otherwise wild type in sequence conferred increases in AAV2 or AAV8 packaging levels.^40, 41^ To decouple whether any potentially improved functions of MAAP variants were due to their novel missense mutations and/or novel C-terminal domains, we included truncation variants L78* and L100* (MAAP-WT2 containing nonsense mutations at the 78^th^ or 100^th^ amino acid positions) as controls (Figure 4C and Table S1). We analyzed the effect of MAAP-SL01 and MAAP-SL08 provided *in trans* on the packaging of recombinant AAV2 encoding GFP. Specifically, recombinant AAV2 packaged by triple transfection of a plasmid encoding the GFP transgene flanked by AAV2 ITRs, pHelper, and pAAV2ι1MAAP in HEK293 cells stably expressing lentivirally-delivered MAAP-SL01 or MAAP-SL08 exhibited statistically significant higher overall titer than production from cells stably expressing MAAP-WT2 (Figure 5A-B). These results indicate that MAAP variants resulting from directed evolution may be useful in industrially relevant recombinant AAV manufacturing processes.

**Figure 5.**
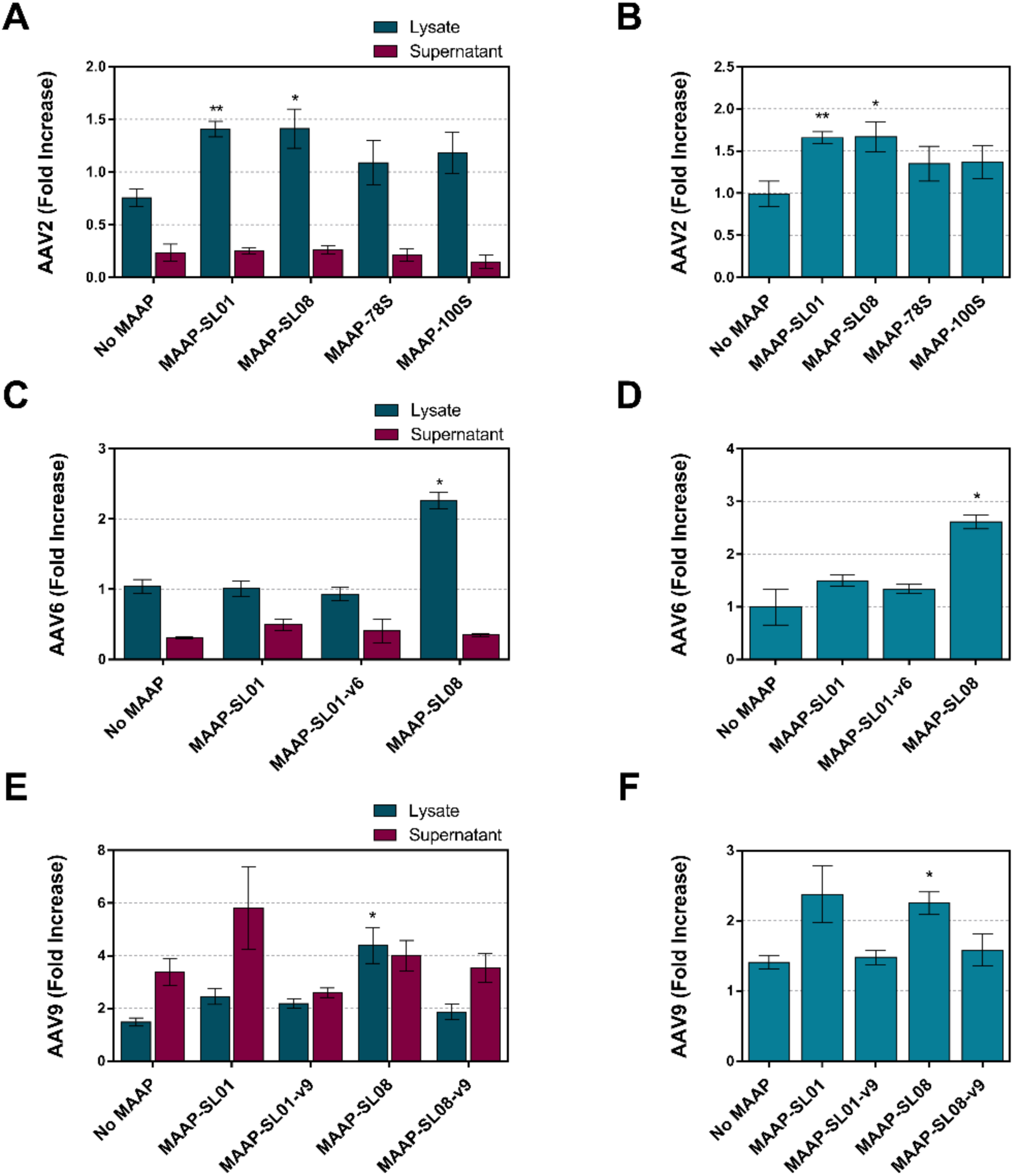
Selected MAAP variants confer increased vector genome packaging in AAV serotypes 2, 6, and 9. A) GFP transgene packaging into AAV2 capsids (encoded by pAAV2Δ MAAP) in the presence of the indicated MAAP variant provided *in trans* via stable expression from the HEK293 cell genome. Both media-associated and cell-associated titers for all conditions were normalized to the cell-associated titer of MAAP-WT2 for each individual biological replicate (N=5 independently packaged biological replicates). B) Total AAV from part A normalized to the total titer of MAAP-WT2 (N=5). C) GFP transgene packaging into AAV6 capsids (encoded by pAAV6ΔMAAP) in the presence of the indicated MAAP variant provided *in trans* via stable expression from the HEK293 cell genome. Both media-associated and cell-associated titers for all conditions were normalized to the cell-associated titer of MAAP-WT6 for each individual biological replicate (N=3). D) Total AAV from part C normalized to the total titer of MAAP-WT6 (N=3). E) GFP transgene packaging into AAV9 capsids (encoded by pAAV9ΔMAAP) in the presence of the indicated MAAP variant provided *in trans* via stable expression from the HEK293 cell genome. Both media-associated and cell-associated titers for all conditions were normalized to the cell-associated titer of MAAP-WT9 for each individual biological replicate (N=3). F) Total AAV from part E normalized to the total titer of MAAP-WT9 (N=3). Error bars, mean ± SEM. *P < 0.05, **P < 0.01 (Student’s t-test).

### MAAP variants isolated from AAV2 screens also enhance recombinant AAV6 and AAV9 packaging

We next characterized whether the evolved MAAP variants showed cross-serotype activity for AAV6 or AAV9 packaging. First, we generated AAV6 (AAV6ι1MAAP) or AAV9 (AAV9ι1MAAP) packaging plasmids for which the endogenous MAAP sequences were ablated by introducing a silent mutation into the near-cognate “CTG” start codon that, like AAV2ι1MAAP, resulted in no change to the amino acid sequence produced by the VP1 ORF. A GFP transgene packaged into recombinant AAV6ι1MAAP or AAV9ι1MAAP by triple transfection in HEK293 cells stably expressing MAAP-SL08 was produced at statistically significant increased levels relative to HEK293 cells stably expressing MAAP-WT6 or MAAP-WT9, respectively (Figure 5C-5F).

## Discussion

Discovered serendipitously as a contaminant of adenovirus preparations in 1965,^42, 43^ AAV is a rare example of a virus that has been extensively studied despite not being pathogenic in any known organism.^1, 44^ Despite AAV’s short genome with a relatively small number of ORFs, many mechanisms underlying its biology are not well understood and, in some cases, completely unknown. For example, the translocation of mature AAV particles away from the nucleus and egress out of the cell require additional investigation. Developing a clear understanding of these processes may improve manufacturing and therapeutic outcomes of AAV-based gene therapies. The discovery and characterization of MAAP illuminates new AAV genomic mutations that may adversely or beneficially affect vector quality and yield. Encoded by an ORF that overlaps with the VP1-encoding ORF, MAAP and the viral VPs naturally co-evolved under strict evolutionary constraints. By uncoupling MAAP from *cap*, such natural evolutionary constraints can be loosened to enable novel mutations, such as SNPs affecting highly conserved VP1 residues or frameshifts resulting in novel C-terminal domains, that would not be possible in a natural evolutionary context.

AAV-based gene therapies currently under clinical evaluation require high doses (∼5 x 10^9^ vector genomes per patient for localized gene therapy applications to ∼1 x 10^17^ vector genomes per patient for applications requiring systemic delivery)^45, 46^ to achieve a therapeutic effect. AAVs are currently utilized in 161 active- or recruiting status clinical trials as of May, 2023,^25^ and robust preclinical research activity feeding the translational pipeline continues.

Furthermore, preclinical and clinical exploration of AAV gene therapies have been expanding from CNS (including retinal) conditions, which in general require lower AAV doses,^47^ to musculoskeletal and other conditions that require high vector doses.^48, 49^ Thus, manufacturing systems face tremendous hurdles to enable AAV gene therapy’s full potential. To achieve sustainable scale-up in the face of increasing demand, innovations leading to higher per-cell output of functional recombinant AAV particles, increased secreted-to-pellet-retained ratios, and lowered full-to-empty and full-to-truncated vector ratios during upstream vector manufacturing processes will be needed.

Here we report a novel directed evolution approach for generating MAAP variants that confer increased quality and quantity of AAV vectors (Figure 3). This approach generated two novel MAAP variants, MAAP-SL01 and MAAP-SL08, that confer increased GFP transgene packaging into AAV2 (Figure 5A-5B). MAAP-SL08 also enabled statistically significant increases in GFP transgene packaging into AAV6 and AAV9, suggesting that MAAP-SL08 acts by mechanisms that are conserved across serotypes (Figure 5C-5F). Although different seeding density, expression level, cell line genotype, and temporal conditions of AAV packaging could conceivably modulate the effects of a given MAAP variant on AAV packaging output, MAAP-SL01 and MAAP-SL08 may have the potential to be directly applied to industrial-scale manufacturing processes. More broadly, our directed evolution approach may be applied to optimize MAAP variants to any given manufacturing process or AAV serotype of interest.

MAAP-SL08 contains unanticipated features that could not have been predicted via rational engineering and whose underlying mechanisms will need to be further characterized. MAAP-SL08 contained the first 26 amino acids of MAAP-WT2 followed by a frameshift into VP1 amino acid positions 54-196 with nine missense mutations scattered three to 20 amino acids apart (Figure 4D). Further work will need to be performed to determine whether improvements conferred by MAAP-SL08 are due to its missense mutations, VP1 fusion region, or a combination thereof. Our second variant, MAAP-SL01, which was the most enriched variant in the dataset, conferred significant increases in AAV2 but not AAV6 or AAV9 packing. These results underscore the possibility that performing directed evolution screens using the serotype of interest as the selective pressure may achieve improved engineering outcomes for those serotypes. MAAP-SL01 contained the first 73 amino acids of MAAP-WT2, with the exceptions of 13 missense mutations, followed by a frameshift mutation that resulted in a novel 26 amino acid-long C-terminal domain (Figure 4D and Table S1). Notably, an “EKE” charge motif located at the far C-terminus of MAAP-SL01 comprised 16.3% of the post-selected dataset. Hypotheses regarding the underlying mechanisms of how individual missense mutations and/or C-terminal charge change affect AAV output deserve further experimental exploration.

Given the marked increase in secretion ratios observed in the bulk selected library population vs wildtype MAAP, it was an unexpected finding that MAAP-SL01 and MAAP-SL08 drove higher overall vector genome titers without significantly increasing the titer of vector genomes secreted to the supernatant. It is possible that packaging AAV2 in the presence of MAAP-SL08 conferred increased specific infectivity of secreted viral particles relative to lower-fitness MAAP variants during packaging selection. Such increase in specific infectivity could be explained by a model for which the MAAP variant, when incorporated into the EV membrane, facilitates increased cellular uptake and/or trafficking of enclosed AAV particles to the nucleus. Thus, the packaging of AAV into EVs and potential effect of MAAP on specific infectivity of such AAV-loaded EVs deserves further investigation. Additional possibilities explaining the striking observed increase in secreted AAV2 vector genomes conferred by the bulk selected MAAP library population relative to MAAP-WT2 (Figure 3F) include the time point at which AAV was sampled following transfection of helper plasmids or that the selected library variants accounting for increase in secreted vector genomes remain unexplored within our dataset. Additional factors worth exploring include how temporal regulation and strength of MAAP expression levels affect AAV packaging. Our stable cell lines expressed MAAP constitutively under a CMV promoter, though it is possible that alternative or inducible promoters may further modulate effects.

In addition to our directed evolution approach and characterization of two leading individual evolved variants, we confirmed prior results, previously shown for AAV8,^41^ that the knockout of MAAP from AAV2 (AAV2Δ1MAAP) results in decreased AAV titer secreted in the supernatant when sampled 72 hours post-transfection (Figure 1A-1F). Our findings that the level of AAV2Δ1MAAP associated with EVs is lower than that of wildtype AAV2, in addition to recent findings that MAAP associates with EVs,^41^ further indicate that MAAP may play a role in incorporating AAV into EVs. MAAP8 was recently found to be important for EV association of AAVs but did not affect the size or number of EVs secreted into the media. Our results in AAV2 closely corroborate this finding. An interesting future avenue would be to determine whether the AAV particles are physically enclosed within EVs or simply tethered to the outer surface of EVs.

More broadly, developing a deeper mechanistic understanding of the pathways and mechanisms that AAVs follow from capsid assembly and genome loading in the nucleus to nuclear escape, intracellular trafficking, and egress will enable engineering of more manufacturable and efficacious AAV gene therapies. The translation of even modest improvements to AAV production in industrial settings can have profound consequences toward reducing global costs of and increasing patient access to these promising, life-saving medicines.

## Materials and Methods

### Plasmids and viruses

#### pSubMAAP

pX601 vector was digested with NcoI + BamHI (New England Biolabs, Ipswich, MA) at 37°C for 60 minutes. The MAAP-WT2 DNA sequence was isolated from the AAV2 genome by Q5 Polymerase-based PCR (New England Biolabs, Ipswich, MA) and cloned into pSubMAAP to assess MAAP effects of MAAP knockout and reconstitution (Figures 1A-F and 2A-F). PCR products were purified using a PCR purification kit (Qiagen, Hilden, Germany) following the manufacturer’s protocol, then digested with NcoI + BamHI at 37°C for 60 minutes. The digested PCR product and pSubMAAP vector was then ligated together using T4 DNA ligase (New England Biolabs, Ipsiwch, MA), transformed into Top10 competent cells (Thermo Fisher Scientific, Waltham, MA), and DNA was isolated and purified by midi prep (Thermo Fisher Scientific, Waltham, MA) following the manufacturer’s protocol. Start codon of MAAP-WT2 (CTG) was substituted with ATG using Q5 Site-Directed Mutagenesis Kit (New England Biolabs, Ipswich, MA) in accordance with manufacturer’s protocol.

#### pMAAP-Library

Error-prone PCR was used to amplify MAAP from the AAV2 genome following the staggered extension process previously described.^50^ Briefly, 10X EP buffer (5 μL), differential volumes of 5 mM MnCl2 (0 μL, 0.2 μL, 0.5 μL, or 0.8 uL, DMSO (2.5 μL), 10X dNTPs (5 μL), 2.5 μL of each 10 μM primer (5’ [AJ005 sequence] 3’ and 5’ [AJ006 sequence] 3’), 1 μL of AAV2 genomes diluted to 1 ng/uL, Taq Polymerase (1 μL), and water (30.5 μL) were mixed together and cycled 33x as shown in Table S3. pSub MAAP was digested with NheI/SpeI in Cutsmart buffer for 1.5 hour at 37C. Ran EP PCR products (inserts) on 1.5% LMT agarose gel. The ∼370 bp band was excised, gel purified using a gel purification kit (Qiagen), digested using NheI/SpeI for four hours, and purified using AMPure XP beads, and ligated with the digested pSubMAAP vector at a 100 ng : 37.5 ng vector : insert ratio using T4 DNA ligase (New England Biolabs, Ipswich, MA). The ligation reactions were desalted for 40 minutes using a membrane on molecular-grade H_2_0 and electroporated into DH10B electrocompetent cells. Electroporated cells were then grown up in overnight cultures and maxi prepped using the manufacturer’s protocol (Thermo Fisher Scientific, Waltham, MA).

#### pAAV2ϕλMAAP

A Q5® Site-Directed Mutagenesis Kit (New England Biolabs, Ipswich, MA) was used to mutate the 19^th^ amino acid position, Leucine (TTG), to a stop codon (TAG) in the MAAP open reading frame, while introducing only a silent mutation in the 45^th^ amino acid position, Leucine (CTT -> CTA) in the VP1 open reading frame of AAV2. Forward (5’GCAGGGGTCTaGTGCTTCCTG3’) and reverse (5’TGTCGTCCTTATGCCGCT3’) primers were used on pRepCap2 (described previously). The plasmid was functionally verified using triple transfection assay in HEK293 cells that successfully packaged AAV2 but produced no MAAP (Figure S1).

#### pAAV9ϕλMAAP

A Q5® Site-Directed Mutagenesis Kit (New England Biolabs, Ipswich, MA) was used to mutate the first amino acid position, Leucine (CTG), to a Proline (CCG) in the MAAP open reading frame, while introducing only a silent mutation in the 27th amino acid position, Proline (CCT -> CCC) in the VP1 open reading frame of AAV9. Unlike the near-cognate start codon CTG, transcriptional initiation off of CCG codons is inefficient and rare in mammalian cells, as repeatedly shown through its seldom occupation in ribosome profiling assays.^51–53^ Primer sequences 5’-CTTTGAAACCcGGAGCCCCTC-3’ and 5’-CCCACCACTCGCGAATTC-3’ were used with pAAV9 as template, and the manufacturer’s protocol was followed.

#### pAAV6ϕλMAAP

In the same manner as pAAV9*ϕλMAAP* generation, Q5® Site-Directed Mutagenesis Kit (New England Biolabs, Ipswich, MA) was used to mutate the first amino acid position, Leucine (CTG), to a Proline (CCG) in the MAAP open reading frame, while introducing only a silent mutation in the 27th amino acid position, Proline (CCT -> CCC) in the VP1 open reading frame of AAV6. Primer sequences 5’-ACTTGAAACCcGGAGCCCCGAAAC-3’ and 5’-CCCACCACTCGCGAATGC-3’ were used with pAAV6 as template, and the manufacturer’s protocol was followed.

#### Lentivirus plasmids for stable cell line generation

pCW57 was digested with ClaI and BamHI. PCR of MAAP inserts was performed using pSubMAAP vectors for which the indicated MAAP variant had previously been incorporated (pSubMAAP-WT2, pSubMAAP-SL01, pSubMAAP-SL08, pSubMAAP-L78*, or pSubMAAP-L100*) as template. The PCR products were digested using ClaI and BamHI in 1X Cutsmart buffer (New England Biolabs, Ipswich, MA) and ligated into the pCW57 backbone using T4 DNA ligase. Samples were transformed into Top10 bacterial cells, and an individual colony was selected and verified using Sanger sequencing.

### Cell lines

#### Parental cells

HEK293 parental cells and HEK293T cells were obtained from the American Type Culture Collection (Manassas, VA, USA) and cultured in DMEM (GIBCO) with 10% fetal bovine serum (Invitrogen) and 1% penicillin/streptomycin/Amphtericin B (GIBCO) at 37 degrees C and 5% CO2.

#### Lentivirus packaging

HEK293T cells (8 x 10^6^ cells per dish) were passaged into 10cm. After16 hrs of incubation, the HEK293T cells were transfected with third-generation lentivirus packaging plasmids using following: DMEM (0.94 mL), lentivirus packaging plasmid with gene of interest (10 ug), pMDLg/pRRE (5 ug), RsRev (2.5 ug), VSV.G (2.5 ug), PEI (60 g). Media was changed 12 hours post-transfection, and HEPES was added at **20mM** final concentration. Following 48 hrs post-transfection, supernatant was harvested in 15ml conical tubes, centrifuged at 3000 rpm for 5 minutes, and filtered using a 200 μM filter syringe unit. The remaining lysate was stored at -80°C for subsequent use.

#### Transduction and selection of cells stably expressing individual MAAP proteins

HEK293 parental cells (which do not contain. T antigen) were plated into three full six-well plates (6 x 10^5^ cells per well) and subsequently infected with packaged lentivirus with polybrene (8 ug/mL). Two days post-transduction, the infection media was replaced with selection media containing puromycin (1 ug/mL). Cells were monitored and passaged until all non-infected cells were dead. Cell lines were used for the first biological replicate of AAV2, AAV6, or AAV9 packaging experiments, and stocks were frozen at -80°C for use in subsequent biological replicates.

### Selection of MAAP variants that confer increased AAV2 production

To select for MAAP variants that confer increase in AAV2 packaging production, the MAAP libraries were first packaged into rAAV2 (pMAAP-Library, pHelper, and pAAV2ΔMAAP) in parental HEK293 cells. At 96 hours post-infection, cell-associated or supernatant-associated AAV was sampled and titered via qPCR using the methods described below. The resulting supernatant-associated MAAP library packaged into AAV2 was then used to infect fresh parental HEK293 cells at MOI = 100, followed by co-transfection of pHelper and pAAV2Δ MAAP at 24 hours post-infection (HPI). AAV was sampled at 96 hours post-infection, and the supernatant-associated AAV was used to infect HEK293 cells at MOI = 100. After four iterations of this process, the total AAV was harvested and prepared for next-generation sequencing.

### Next-generation sequencing and bioinformatics

Next-generation sequencing of the pre- and post-selected MAAP library was performed to evaluate the fold enrichment of each MAAP sequence following execution of selections for AAV packaging and secretion to the cell culture media. The MAAP sequence was amplified (forward primer: 5’ GCGCctggctaactaccggtgctagc-3’ and reverse primer: 5’-GCCAgtaatctggaacatcgtatgggtagActaGt -3’) from the AAV genomes following the fourth round of selection by PCR using Q5 High-Fidelity DNA polymerase (New England Biolabs, Ipswich, MA) and 21 cycles of PCR. PCR products were purified using AMPure XP Beads (Beckman Coulter, Brea, CA) using the manufacturer’s protocol. The resulting product was quantified using a Qubit dsDNA HS assay kit and Qubit fluorometer (Thermo Fisher Scientific, Waltham, MA), and a total of 30 μg was used as template for 10 PCR cycles using IDT-8 TruSeq primers for barcoding. The barcoded amplicons were then pooled and sequenced at the QB3 Vincent J. Coates Genomics Sequencing Laboratory at UC Berkeley using the 300 PE MiSeq platform and TruSeq SBS Kit v3-HS (Illumina, San Diego, CA). Illumina FASTQ files were demultiplexed, and adaptor sequences were trimmed. The resulting demultiplexed sequences were analyzed using Geneious software (Dotmatics, Boston, MA) to determine the identity and read count of each MAAP sequence.

### AAV packaging to assess MAAP variant function

Stable cell lines expressing selected variants of MAAP were triple transfected using PEI with pHelper, AAV-Rep/Cap ΔMAAP, and the ss-CAG-GFP. At 4-days post transfection, supernatant and cells were harvested to determine AAV titer using detergent-based method as previously described.^54^

### qPCR

To assess AAV titer in the supernatant, cell supernatant was collected at the indicated time point. Cell debris was removed by centrifugation at 3000 rpm in a table-top centrifuge for 5 minutes. Next, MgCl_2_ and Benzonase (Millipore, Hayward, CA) were added per mL of supernatant sample and incubated for 1hr at 37°C. After one hour, Benzonase was neutralized by adding NaCl per mL of sample. Samples were then ten-fold diluted with DI water and assayed by qPCR to quantify amounts of GFP transgene as described below. To assess AAV titer associated with the cell pellet, cell media was replaced with 4mL of DMEM containing 10% Triton X-100, Benzonase, and MgCl_2_. Cells were then incubated for 1 hour at 37°C and were rocked every 15 minutes. After 1hr, 1mL of cell media containing detached cells was collected and diluted ten-fold with DI water for subsequent qPCR analysis. qPCR was used to quantify the titer of AAV particles that contain genomes using SYBR Green, 3 mM MgCl2, 0.2 mM dNTP, and Jump Start Taq on a CFX RT PCR machine (Bio-Rad, Hercules, CA). Plasmid DNA was used from a concentration of 1 ng/μL to 0.0001 ng/μL to generate a standard curve with primers against CMV (5’-ATGGTGATGCGGTTTTGGCAG-3’ and 5’-GGCGGAGTTGTTACGACATTTTGG-3’) or GFP (5’-ACTACAACAGCCACAACGTCTA TATCA-3’ and 5’ -GGCGGATCTTGAAGTTCACC-3’). Each biological replicate represented a separately transfected T-25 flask, where the number of plates ranged from n=3 to n=6 performed on separate days. Each biological replicate was assayed with two technical replicates during qPCR analysis.

### Capsid quantification

Fully assembled capsids of AAV2 were quantified using sandwich ELISA (PROGEN, AAV Titration ELISA 2.0R) according to manufacturer’s protocol. Each sample from the lysate or supernatant was diluted 200-fold and analyzed using ELISA reader at 450 nm wavelength (Bio-Tek uQUANT Microplate Spectrophotometer). Viral genome copy number of each sample was normalized to capsid quantification to calculate the full and empty capsid ratio.

### Immunoblot analysis

HEK293T cells were transfected with pSubMAAP using PEI. At 36-hours post-transfection, cells were harvested and lysed with RIPA buffer (20 mM Tris-HCl, pH 7.4, 150 mM NaCl, 1% NP-40, and 1 mM EDTA) containing a protease inhibitor cocktail (Thermo Fisher Scientific, Waltham, MA). Whole cell lysates were loaded into pre-prepared SDS-PAGE gel (Invitrogen, Waltham, MA) and transferred to PVDF membrane. Anti-HA antibody (Cell Signaling, #3724) (1:1000) and HRP-conjugated anti-Rabbit IgG antibody (Cell Signaling, #7074) (1:3000) were used for immunoblotting.

### Extracellular vesicle isolation and quantification

We harvested supernatant from HEK293T cells grown in Dulbecco’s Modified Eagle’s Medium (DMEM) supplemented with 10% fetal bovine serum (FBS) and conditioned to remove pre-existing EVs in seven 15 cm cell culture dishes (Thermo Fisher Scientific, Waltham, MA). All subsequent manipulations were completed at 4°C. Cells and large debris were removed by low-speed sedimentation at 1000 × g for 15 minutes followed by medium-speed sedimentation at 10,000 × g for 15 minutes in an F10-6X500Y rotor (Sorvall). EVs were pelleted from conditioned media by spinning at ∼100,000 x g (29,500 rpm) for one hour in a SW32 Ti rotor (Beckman), and the supernatant was discarded. Crude EVs were resuspended in PBS. Resuspended crude EVs were spun at 1000 × g for 5 minutes to remove any aggregates formed during centrifugation. A solution of 65% Sucrose, 20 mM Tris pH 7.4 were added to resuspended crude EVs until the solution reached 55% sucrose, and 3 mL of this solution was added to an SW55 tube. Next, 1 mL of 40% sucrose, 20 mM Tris pH 7.4 followed by one mL of 10% sucrose, 20 mM Tris pH 7.4 was carefully layered onto the gradient and centrifuged in a SW55 rotor (Beckman) at 36,500 rpm for 16 hours with minimum acceleration and no brake. Fractions (400 µl) were collected from top to bottom and analyzed by Western Blot. Density measurements were taken using a refractometer.

### Statistical analysis

Raw data were recorded electronically and statistical analyses were performed with Microsoft Excel or Prism software version 9 (GraphPad Software, San Diego, CA). Vector titers were expressed as means ± SEM with Student’s *t*-test. An α value (*P*) of 0.05 was considered as statistically significant, and all *P* values were two sided.

## Acknowledgments

We acknowledge Prof. Randy Schekman for sharing his helpful insights on EVs and Juan Hurtado for his consultation regarding next-generation sequencing analysis.

## Author Contributions

AJS co-conceived of the study design, contributed to all performed experiments, analyzed data, and wrote manuscript. HL co-conceived of study design, contributed to all performed experiments, analyzed data, and edited manuscript. VK and AC contributed to cloning, stable cell line generation and maintenance, and assessment of infectious titer. JKW performed EV sampling and analysis depicted in Figure 2A-2F. DVS supervised the project, co-conceived study design, and edited manuscript.

