## Supplemental Information for "Evolving Membrane-Associated Accessory Protein Variants for Improved Adeno-associated Virus Production"

**This PDF file includes:**

Supplemental Figures S1-S3

Supplemental Tables S1-S3

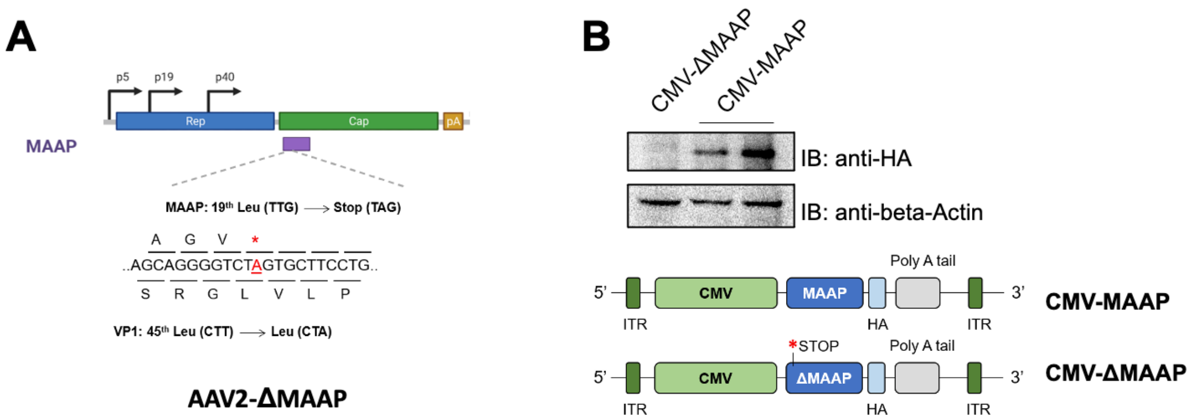


**Figure S1. Approach to study MAAP function in AAV2.** A) Endogenous MAAP expression from the AAV2 *cap* region was eliminated by the introduction of an early stop codon at the 19$^{th}$ Leucine position in MAAP without affecting the translated amino acid sequence of the VP1 gene product. B) Western blot analysis confirming the knockout of MAAP expression.

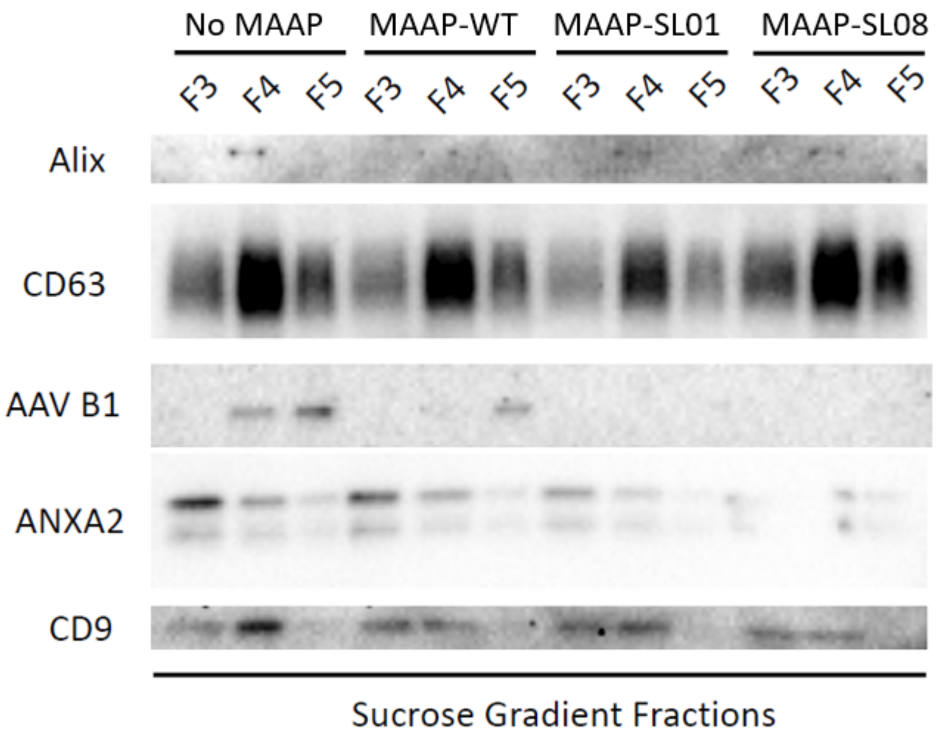


**Figure S2. Western blot of sucrose gradient fractions.** AAV-GFP was packaged with AAV2-WT or AAV2-MAAP-null packaging plasmids in HEK293 cells stably expressing MAAP-WT, MAAP-SL01, or MAAP-SL08. Cell culture media was harvested at 3-day post transfection for exosome analysis as described in Figure 2. Each fraction was immunoblotted using antibodies targeting exosome markers or AAV capsid protein.

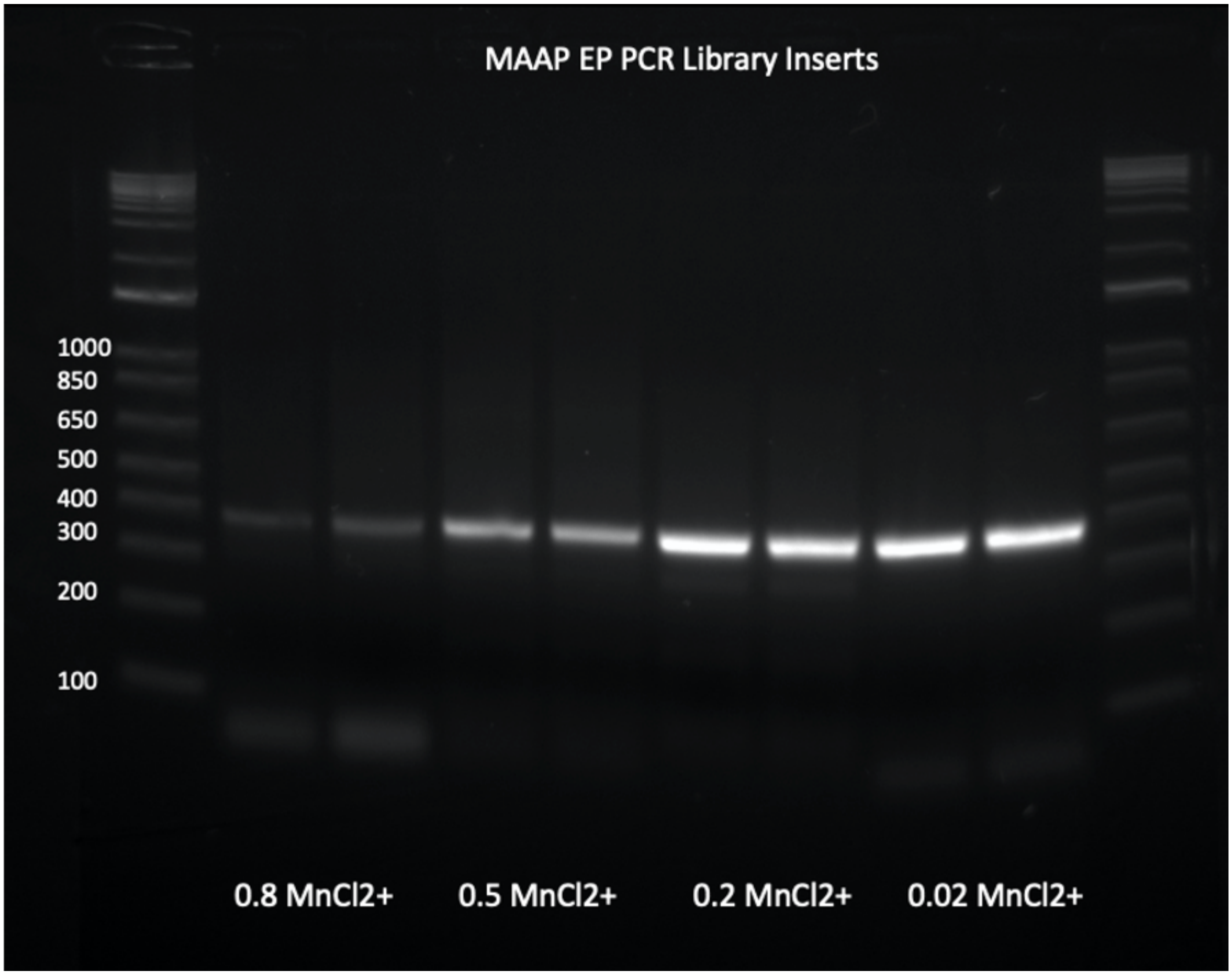


**Figure S3. Generation of MAAP library using error-prone PCR.** Gel electrophoresis of the MAAP coding region amplified by error-prone PCR using different (0.8 μL, 0.5μL, 0.2 μL, or 0.02 μL) volumes of 5 mM $MnCl2^+$ in 25 μL reactions. The concentration of $MnCl2^+$ is inversely related to the fidelity of the T4 DNA polymerase, resulting high mutation rate and lowered full-length PCR products at high concentrations.

**Supplemental Table 1**

| | |
|---|---|
| MAAP-WT2 | MAHHHQSPQSGIRTTAGVLCFLGTSTSDPSTDSTRESRSTRQTPRPS<br>STTKPTTGSSTAETTRTSSTTTPTRSFRSALKKIRLLGATSDEQSSRR<br>KRGFLNLWAWLRNLLRRLREKRGR* |
| MAAP-WT6 | MEPRNPKPTSKSRTTAGVWCFLATSTSDPSTDSTRGSPSTRRMQRPS<br>STTRPTTSSSKRVTIRT<br>CGITTPTPSFRSVCKKIRLLGATSGEQSSRPRRGFSNLLVWLRKVLR<br>RLLERNVR* |
| MAAP-WT9 | MEPLNPRQINNIKTTLEVLCFRVTNTLDPATDSTRGSRSTQQTRRPSS<br>TTRPTTSSSRPETTRT<br>SSTTTPTPSSRSGSKKIRLLGATSGEQSSRPKRGFLNLLVWLRKRLR<br>RLLERRGL* |
| MAAP-L78* | MAHHHQSPQSGIRTTAGVLCFLGTSTSDPSTDSTRESRSTRQTPRPS<br>STTKPTTGSSTAETTRTSSTTTPTRSFRSA* |
| MAAP-L100* | MAHHHQSPQSGIRTTAGVLCFLGTSTSDPSTDSTRESRSTRQTPRPS<br>STTKPTTGSSTAETTRTSSTTTPTRSFRSALKKIRLLGATSDEQSSRR<br>KRGF* |
| MAAP-SL01 | MAHHHQGPQRGIRATAGVLCFLGTSTSDPSTDTSRESRSTRQTPRPL<br>SMSKPTTGSPTADNTRTSSATSPTRSLGAPKGCYVFRGQTRTSSLPG<br>EKEGS* |
| MAAP-SL08 | MAHHHQSPQSGIRTTAGVLCFLGTSTGPFSGLDKGEPVIEADAAAL<br>DHDKAYDRQLDGGDNLYLKYNHADAEFQERLKEDTSCGGNLGRA<br>VFQANKRVPEPLGLVEKPVKTAPGKKRPG* |

| | |
|---|---|
| MAAP-SL01V6 | MEPRNPKPTSKSRTTAGVWCFLATSTSDPSTDSTRGSPSTRRMQRPS<br>STTRPTTSSSKRVTIRT<br>CGITTPTPSFRSVCKKIRLLGATRTSSLPGEKEGS* |
| MAAP-SL01V9 | MEPLNPRQINNIKTTLEVLCFRVTNTLDPATDSTRGSRSTQQTRRPSS<br>TTRPTTSSSRPETTRT<br>SSTTTPTPSSRSGSKKIRLLGATRTSSLPGEKEGS* |
| MAAP-9VPF | LEPLNPRQINNIKTTLEVLCFRVTNTLDPATDSTRGSRSTQQTRRPSST<br>TRPTTSSSRPETTRTSSTTTPTPSSRSGSKKIRLLGATSGEQSSRPKRGF<br>LPLGLVEEAAKTAPGKKRPVEQSPQEPDSSAGIGKSGAQPAKKRLNF<br>GQTGDTESVPDPQPIGEPPAAPSGVGSLTMASGGGAPVADNNEGAD<br>GVGSSSGNWHCDSQWLGDRVITTSTRTWALPTYNNHLYKQISNSTS<br>GGSSNDNAYFGYSTPWGYFDFNRFHCHFSPRDWQRLINNNWGFRP<br>KRLNFKLFNIQVKEVTDNNGVKTIANNLTSTVQVFTDSDYQLPYVL<br>GSAHEGCLPPFPADVFMIPQYGYLTLNDGSQAVGRSSFYCLEYFPSQ<br>MLRTGNNFQFSYEFENVPFHSSYAHSQSLDRLMNPLIDQYLYYLSKT<br>INGSGQNQQTLKFSVAGPSNMAVQGRNYIPGPSYRQQRVSTTVTQN<br>NNSEFAWPGASSWALNGRNSLMNPGPAMASHKEGEDRFFPLSGSLIF<br>GKQGTGRDNVDADKVMITNEEEIKTTNPVATESYGQVATNHQSAQA<br>QAQTGWVQNQGILPGMVWQDRDVYLQGPIWAKIPHTDGNFHPSPL<br>MGGFGMKHPPPQILIKNTPVPADPPTAFNKDKLNSFITQYSTGQVSV<br>EIEWELQKENSKRWNPEIQYTSNYYKSNNVEFAVNTEGVYSEPRPIG<br>TRY LTRNL* |

Supplemental Table 1: Amino acid sequences of MAAP variants used in this study.

**Supplemental Table 2**

| | |
|---|---|
| MAAP-WT2 | ATGGCCCACCACCACCAAAGCCCGCAGAGCGGCATAAGGACGA<br>CAGCAGGGGTCTTGTGCTTCCTGGGTACAAGTACCTCGGACCCT<br>TCAACGGACTCGACAAGGGAGAGCCGGTCAACGAGGCAGACGC<br>CGCGGCCCTCGAGCACGACAAAGCCTACGACCGGCAGCTCGAC<br>AGCGGAGACAACCCGTACCTCAAGTACAACCACGCCGACGCGG<br>AGTTTCAGGAGCGCCTTAAAGAAGATACGTCTTTTGGGGGCAAC<br>CTCGGACGAGCAGTCTTCCAGGCGAAAAAGAGGGTTCT<br>TGAACCTCTGGGCCTGGTTGAGGAACCTGTTAAGACGGCTCCGG<br>GAAAAAAGAGGCCGGTAA |
| MAAP-WT6 | ATGGAGCCCCGAAACCCAAAGCCAACCAGCAAAAGCAGGACGA<br>CGGCCGGGGTCTGGTGCTTCCTGGCTACAAGTACCTCGGACCCT<br>TCAACGGACTCGACAAGGGGGAGCCCGTCAACGCGGCGGATGC<br>AGCGGCCCTCGAGCACGACAAGGCCTACGACCAGCAGCTCAAA<br>GCGGGTGACAATCCGTACCTGCGGTATAACCACGCCGACGCCGA<br>GTTTCAGGAGCGTCTGCAAGAAGATACGTCTTTTGGGGGCAACC<br>TCGGGCGAGCAGTCTTCCAGGCCAAGAAGAGGGTTCTCGAACC<br>TTTTGGTCTGGTTGAGGAAGGTGCTAAGACGGCTCCTGGAAAGA<br>AACGTCCGGTAG |
| MAAP-WT9 | ATGGAGCCCCTCAACCCAAGGCAAATCAACAACATCAAGACAAC<br>GCTCGAGGTCTTGTGCTTCCGGGTTACAAATACCTTGGACCCGGC<br>AACGGACTCGACAAGGGGGAGCCGGTCAACGCAGCAGACGCGG<br>CGGCCCTCGAGCACGACAAGGCCTACGACCAGCAGCTCAAGGC |

| | |
|---|---|
| | CGGAGACAACCCGTACCTCAAGTACAACCACGCCGACGCCGAG<br>TTCCAGGAGCGGCTCAAAGAAGATACGTCTTTTGGGGGCAACCT<br>CGGGCGAGCAGTCTTCCAGGCCAAAAAGAGGCTTCTTGAACCTC<br>TTGGTCTGGTTGAGGAAGCGGCTAAGACGGCTCCTGGAAAGAA<br>GAGGCCTGTAG |
| MAAP-L78* | ATGGCCCACCACCACCAAAGCCCGCAGAGCGGCATAAGGACGA<br>CAGCAGGGGTCTTGTGCTTCCTGGGTACAAGTACCTCGGACCCT<br>TCAACGGACTCGACAAGGGAGAGCCGGTCAACGAGGCAGACGC<br>CGCGGCCCTCGAGCACGACAAAGCCTACGACCGGCAGCTCGAC<br>AGCGGAGACAACCCGTACCTCAAGTACAACCACGCCGACGCGG<br>AGTTTCAGGAGCGCCTAA |
| MAAP-L100* | ATGGCCCACCACCACCAAAGCCCGCAGAGCGGCATAAGGACGA<br>CAGCAGGGGTCTTGTGCTTCCTGGGTACAAGTACCTCGGACCCT<br>TCAACGGACTCGACAAGGGAGAGCCGGTCAACGAGGCAGACGC<br>CGCGGCCCTCGAGCACGACAAAGCCTACGACCGGCAGCTCGAC<br>AGCGGAGACAACCCGTACCTCAAGTACAACCACGCCGACGCGG<br>AGTTTCAGGAGCGCCTTAAAGAAGATACGTCTTTTGGGGGCAAC<br>CTCGGACGAGCAGTCTTCCAGGCGAAAAAGAGGGTTCTAA |
| MAAP-SL01 | ATGGCCCACCACCACCAAGGCCCGCAGAGGGGCATAAGGGCCA<br>CAGCAGGGGTCTTGTGCTTCCTGGGTACAAGTACCTCGGACCCT<br>TCAACGGACACCAGCAGGGAGAGCCGGTCAACGAGGCAGACGC<br>CGCGGCCGCTGAGCATGAGCAAGCCTACGACCGGCAGCCCGAC<br>AGCGGATAACACCCGTACCTCAAGTGCGACCAGCCCGACGCGGA |

| | |
|---|---|
| | GTCTGGGCGCGCCGAAAGGCTGCTATGTGTTTCGCGGCCAGACC<br>CGCACCAGCAGCCTGCCGGGCGAAAAAGAAGGCAGCTAA |
| MAAP-SL08 | ATGGCCCACCACCACCAAAGCCCGCAGAGCGGCATAAGGACGA<br>CAGCAGGGGTCTTGTGCTTCCTGGGTACAAGTACCGGACCCTTC<br>AGCGGACTCGACAAGGGAGAGCCGGTCATTGAGGCAGACGCCG<br>CGGCCCTCGATCACGACAAAGCCTACGACCGGCAGCTCGACGGT<br>GGAGACAACCTGTACCTCAAGTACAACCACGCCGACGCGGAGTT<br>TCAGGAGCGCCTTAAAGAAGATACGTCTTGTGGGGGCAACCTCG<br>GACGAGCAGTCTTCCAGGCGAATAAGAGGGTTCCGGAACCTCTG<br>GGCCTGGTTGAGAAACCTGTTAAGACGGCTCCGGGAAAAAAGA<br>GGCCGGGCTAA |
| MAAP-SL01V6 | ATGGAGCCCCGAAACCCAAAGCCAACCAGCAAAAGCAGGACGA<br>CGGCCGGGGTCTGGTGCTTCCTGGCTACAAGTACCTCGGACCCT<br>TCAACGGACTCGACAAGGGGGAGCCCGTCAACGCGGCGGATGC<br>AGCGGCCCTCGAGCACGACAAGGCCTACGACCAGCAGCTCAAA<br>GCGGGTGACAATCCGTACCTGCGGTATAACCACGCCGACGCCGA<br>GTTTCAGGAGCGTCTGCAAGAAGATACGTCTTTTGGGGGCAACC<br>CGCACCAGCAGCCTGCCGGGCGAAAAAGAAGGCAGCTAA |
| MAAP-SL01V9 | ATGGAGCCCCTCAACCCAAGGCAAATCAACAACATCAAGACAAC<br>GCTCGAGGTCTTGTGCTTCCGGGTTACAAATACCTTGGACCCGGC<br>AACGGACTCGACAAGGGGGAGCCGGTCAACGCAGCAGACGCGG<br>CGGCCCTCGAGCACGACAAGGCCTACGACCAGCAGCTCAAGGC<br>CGGAGACAACCCGTACCTCAAGTACAACCACGCCGACGCCGAG |

| | |
|---|---|
| | TTCCAGGAGCGGCTCAAAGAAGATACGTCTTTTGGGGGCAACCC GCACCAGCAGCCTGCCGGGCGAAAAAGAAGGCAGCTAA |
| MAAP-9VPF | CTGGAGCCCCTCAACCCAAGGCAAATCAACAACATCAAGACAA CGCTCGAGGTCTTGTGCTTCCGGGTTACAAATACCTTGGACCCGG CAACGGACTCGACAAGGGGGAGCCGGTCAACGCAGCAGACGCG GCGGCCCTCGAGCACGACAAGGCCTACGACCAGCAGCTCAAGG CCGGAGACAACCCGTACCTCAAGTACAACCACGCCGACGCCGA GTTCCAGGAGCGGCTCAAAGAAGATACGTCTTTTGGGGGCAACC TCGGGCGAGCAGTCTTCCAGGCCAAAAAGAGGCTTCTTGCCTCT TGGTCTGGTTGAGGAAGCGGCTAAGACGGCTCCTGGAAAGAAG AGGCCTGTAGAGCAGTCTCCTCAGGAACCGGACTCCTCCGCGGG TATTGGCAAATCGGGTGCACAGCCCGCTAAAAAGAGACTCAATT TCGGTCAGACTGGCGACACAGAGTCAGTCCCAGACCCTCAACC AATCGGAGAACCTCCCGCAGCCCCCTCAGGTGTGGGATCTCTTA CAATGGCTTCAGGTGGTGGCGCACCAGTGGCAGACAATAACGAA GGTGCCGATGGAGTGGGTAGTTCCTCGGGAAATTGGCATTGCGA TTCCCAATGGCTGGGGGACAGAGTCATCACCACCAGCACCCGAA CCTGGGCCCTGCCCACCTACAACAATCACCTCTACAAGCAAATCT CCAACAGCACATCTGGAGGATCTTCAAATGACAACGCCTACTTC GGCTACAGCACCCCCTGGGGGTATTTTGACTTCAACAGATTCCAC TGCCACTTCTCACCACGTGACTGGCAGCGACTCATCAACAACAA CTGGGGATTCCGGCCTAAGCGACTCAACTTCAAGCTCTTCAACAT TCAGGTCAAAGAGGTTACGGACAACAATGGAGTCAAGACCATCG |

| | |
|---|---|
| | CCAATAACCTTACCAGCACGGTCCAGGTCTTCACGGACTCAGAC TATCAGCTCCCGTACGTGCTCGGGTCGGCTCACGAGGGCTGCCT CCCGCCGTTCCCAGCGGACGTTTTCATGATTCCTCAGTACGGGTA TCTGACGCTTAATGATGGAAGCCAGGCCGTGGGTCGTTCGTCCTT TTACTGCCTGGAATATTTCCCGTCGCAAATGCTAAGAACGGGTAA CAACTTCCAGTTCAGCTACGAGTTTGAGAACGTACCTTTCCATAG CAGCTACGCTCACAGCCAAAGCCTGGACCGACTAATGAATCCAC TCATCGACCAATACTTGTACTATCTCTCAAAGACTATTAACGGTTC TGGACAGAATCAACAAACGCTAAAATTCAGTGTGGCCGGACCCA GCAACATGGCTGTCCAGGGAAGAAACTACATACCTGGACCCAGC TACCGACAACAACGTGTCTCAACCACTGTGACTCAAAACAACAA CAGCGAATTTGCTTGGCCTGGAGCTTCTTCTTGGGCTCTCAATGG ACGTAATAGCTTGATGAATCCTGGACCTGCTATGGCCAGCCACAA AGAAGGAGAGGACCGTTTCTTTCCTTTGTCTGGATCTTTAATTTT TGGCAAACAAGGAACTGGAAGAGACAACGTGGATGCGGACAAA GTCATGATAACCAACGAAGAAGAAATTAAAACTACTAACCCGGT AGCAACGGAGTCCTATGGACAAGTGGCCACAAACCACCAGAGT GCCCAAGCACAGGCGCAGACCGGCTGGGTTCAAAACCAAGGAA TACTTCCGGGTATGGTTTGGCAGGACAGAGATGTGTACCTGCAA GGACCCATTTGGGCCAAAATTCCTCACACGGACGGCAACTTTCA CCCTTCTCCGCTGATGGGAGGGTTTGGAATGAAGCACCCGCCTC CTCAGATCCTCATCAAAAACACACCTGTACCTGCGGATCCTCCAA CGGCCTTCAACAAGGACAAGCTGAACTCTTTCATCACCCAGTATT |

| | |
|---|---|
| | CTACTGGCCAAGTCAGCGTGGAGATCGAGTGGGAGCTGCAGAA GGAAAACAGCAAGCGCTGGAACCCGGAGATCCAGTACACTTCC AACTATTACAAGTCTAATAATGTTGAATTTGCTGTTAATACTGAAG GTGTATATAGTGAACCCCGCCCCATTGGCACCAGATACCTGACTC GTAATCTGTAA |

Supplemental Table 2: Nucleic acid sequences of MAAP variants used in this study.

**Supplemental Table 3**

| Cycle | Step | Number of Repeats | Temperature | Time (minutes:seconds) |
|---|---|---|---|---|
| 1 | 1 | 1X | 95°C | 5:00 |
| 2 | 1 | 33X | 95°C | 0:30 |
| | 2 | | 56°C | 0:30 |
| | 3 | | 68°C | 1:00 |
| 3 | 1 | 1X | 68°C | 10:00 |
| 4 | 1 | NA | 4°C | Hold |

Supplemental Table 3: Thermocycler conditions for error-prone PCR to generate MAAP library
